# Selection in response to community diversity alters plant performance and functional traits

**DOI:** 10.1101/158709

**Authors:** Sofia J. van Moorsel, Marc W. Schmid, Terhi Hahl, Debra Zuppinger-Dingley, Bernhard Schmid

## Abstract

In grassland biodiversity experiments the positive biodiversity-ecosystem functioning relationship generally increases over time. However, we know little about the underlying short-term evolutionary processes. Using five plant species selected for twelve years in a biodiversity experiment in mixture or monoculture and plants without such a selection history, we assessed whether differential selection altered productivity, biodiversity effects, and functional trait differences within newly assembled monocultures and 2-species mixtures. Plants without past community selection history produced the lowest assemblage biomass and showed the weakest biodiversity effects. In newly assembled mixtures, plants with a selection history in mixtures produced more biomass than plants with a monoculture selection history. Biodiversity effects were generally positive and differed significantly between selection histories. However, contrary to our expectations, biodiversity effects were not stronger for mixture-type plants. Biodiversity effects were influenced by both trait differences between plants and community-weighted means, but these relationships were mostly independent of selection history. Our findings suggest that twelve years of selection history in monocultures or species mixtures differentiated plants of each species into monoculture- and mixture-types. Such rapid evolution of different community-types within grassland species and its effect on ecosystem services and functioning are likely to be important for species conservation practice.

## 1. Introduction

The importance of biodiversity for maintaining ecosystem functions such as productivity or nutrient cycling is now well established (Cardinale et al., 2012). The biodiversity-productivity relationship is positive in grassland ecosystems (e.g. Isbell et al., 2011; Tilman et al., 2001), with biodiversity increasing multiple ecosystem functions (Soliveres et al., 2016). The positive effect of biodiversity strengthens with time (Cardinale et al., 2007; Reich et al., 2012), suggesting that complementarity between the co-occurring species can increase over time (Fargione et al., 2007).

Despite more than a decade of research on the biodiversity-productivity relationship (e.g. Reich et al., 2012), we know little about the evolutionary mechanisms that potentially affect species interactions (Thorpe et al., 2011). Selection acting on traits may increase ecological combining ability (Aarssen, 1983; Harper, 1977) via niche differentiation in plant mixtures (Zuppinger-Dingley et al., 2014). Such adaptation occurs when there is either sufficient standing genetic variation in a population and a sorting out of the most suitable genotypes (Fakheran et al., 2010) or by recombination and novel mutations (Anderson et al., 2011). Furthermore, plants may adapt to a novel environment by phenotypic plasticity (Price et al., 2003; Turcotte and Levine, 2016), thus changing their morphology without genotypic changes. Epigenetic variation may contribute to phenotypic variation and thus provides a further source for selection and adaptation (Bossdorf et al., 2008).

Whereas the influence of environmental factors on adaptive responses of plant populations is well studied (e.g., Joshi et al., 2001; Schmid, 1985), much less effort has been devoted to studying the influence of community diversity on a species’ performance (but see Kleynhans et al., 2016; Lipowsky et al., 2011). Based on previous observations in experimental ecosystems suggesting a “division of labor” among species in plant mixtures, it is likely that community diversity plays a role in the evolution of plant functional trait variation. For example, in forests more diverse tree communities express greater crown complementarity (Niklaus et al., 2017; Williams et al., 2017). In diverse grassland communities increased complementarity effects, as estimated by the additive partitioning method of Loreau and Hector (2001), promote community productivity via diversification of the canopy structure and hence light and space use (Allan et al., 2011; Spehn et al., 2000), soil resource partitioning (Fornara and Tilman, 2008; Roscher et al., 2008; von Felten et al., 2009), root depth distribution (Mueller et al., 2013) and distribution of leaf mass (Wacker et al., 2009). However, these *in situ* observations do not allow separating the contribution of phenotypic plasticity from potential underlying shifts in the population structure due to selection of different community-types. To understand to which extent evolutionary processes drive these differences, it is thus important to assess biodiversity effects in a common environment.

Using the additive partitioning method (Loreau and Hector, 2001), the net biodiversity effect (NE) can be partitioned into complementarity (CE) and sampling effects (SE). When the CE drives over-yielding, most species are expected to contribute to higher productivity in more diverse communities. In contrast, when the SE drives over-yielding, a few dominant species increase community productivity in species mixtures. The CE is therefore related to coexistence and trait variation between species as it inherently suggests a differentiation in functional traits (Cadotte et al., 2009; Flynn et al., 2011). Conversely, the SE should be driven by traits of the dominant species and thus by community-weighted trait means (CWMs); an increase in CWMs (e.g., taller plants) should increase biodiversity effects (Cadotte, 2017; Roscher et al., 2012).

The use of functional traits to define species’ niches has a long history in evolutionary ecology (Roughgarden, 1974; Schoener and Gorman, 1968; van Valen, 1965). Recently it has become a popular approach in functional ecology (Violle et al., 2007) to explain mechanisms of species coexistence and ecosystem functioning (Hart et al., 2016; Kraft et al., 2015). However, we do not know how such trait-based niches and the associated functional traits may evolve (Roscher et al., 2015; Sterck et al., 2011).

We tested whether community diversity can act as a selective environment shaping biodiversity effects and functional traits. To test for the heritability of these effects and traits, we grew offspring of plants grown for twelve years in a biodiversity experiment in monocultures and two-species mixtures in a common environment. We measured biomass production and traits of individual plants in monocultures and mixtures established with seedlings from either a selection history of experimental monoculture or mixture communities in a biodiversity field experiment (Jena Experiment, see Roscher et al., (2004) for details) or in monoculture fields from the commercial seed supplier which provided the original seeds for the Jena Experiment in 2002. We refer to the plants growing in Jena since 2002 in mixture or monoculture as mixture- and monoculture-type plants, respectively. The plants derived from seeds obtained from the commercial supplier in 2014 are referred to as naïve plants.

We previously assessed selection outcomes in the Jena Experiment after eight years and one controlled sexual reproduction cycle (Zuppinger-Dingley et al., 2014). Here, we prolonged the selection treatment by four more years, added a second controlled sexual reproduction cycle, and refined our insights by measuring all individuals in test communities, thus also allowing us to assess intra-specific variation within communities. We included naïve plants as a control treatment without selection. We hypothesized that during the twelve years of selection in the experimental field, mixture-type plants may have evolved increased mixture performance. In turn, this may be associated with a larger NE, in particular CE, and larger between-species trait variations. Conversely, we expected monoculture-type plants to have evolved increased monoculture performance, which should be related to larger within-species trait variation.

## 2. Materials and methods

### 2.1 Plant selection histories

To test whether plant types selected over twelve years in mixtures outperform those types selected in monocultures when assembled in mixture test communities, we chose five species grown in monoculture and mixture plots in the Jena Experiment (Jena, Thuringia, Germany, 51°N, 11°E, 135 m a.s.l., see Roscher et al., (2004) for experimental details): *Plantago lanceolata* L.*, Prunella vulgaris* L.*, Veronica chamaedrys* L., *Galium mollugo* L. and *Lathyrus pratensis* L. The study species had previously been classified into the following functional groups (Roscher et al. 2004): small herbs *(V. chamaedrys, P. vulgaris,* and *P. lanceolate)*, tall herb *(G. mollugo)* and legumes (*L. pratensis)*. Plants with Jena Experiment selection history were sown in either mixture or monoculture in 2002 (selection history “mixture” and “monoculture”, respectively) and had undergone twelve years of selection from 2002 until 2014 in either plant monocultures or species mixtures. The species compositions in the experimental plots in Jena were maintained by weeding three times per year in spring, summer and autumn and by mowing twice per year at peak biomass in spring and summer.

We used plant progeny from three different selection histories for the experiment. Plants without selection history in the Jena Experiment (selection history “naïve”) were obtained from a commercial seed supplier (Rieger Hoffmann GmbH, Blaufelden-Raboldshausen, Germany).

### 2.2 First controlled seed production

In spring 2010, plant communities of 48 plots (12 monocultures, 12 two-species mixtures, 12 four-species mixtures and 12 eight-species mixtures) of the Jena Experiment (Roscher et al., 2004) were collected as cuttings and transplanted into an experimental garden in slug-exclosure compartments at the University of Zurich, Switzerland Switzerland (47°33'N, °37 'E, 534 m a.s.l.), in the identical plant composition as the original experimental plots for the first controlled sexual reproduction among co-selected plants (Zuppinger-Dingley et al., 2014). In spring 2011, the seedlings produced from the seeds of the first controlled sexual reproduction in Zurich were transplanted back into those plots of the Jena Experiment from where the parents had originally been collected. In these newly established plots, plant communities with an identical composition to the original communities were maintained for three years until 2014.

### 2.3 Second controlled seed production

To ensure a second sexual reproductive event for the collection of seed material, entire plant communities from the experimental plots replanted in Jena in 2011 were excavated in March 2014. As for the first controlled seed production, the plants from Jena were used to establish plots with an identical plant composition in the experimental garden at the University of Zurich,. We added a 30 cm layer of soil (1:1 mixture of garden compost and field soil, pH 7.4, commercial name Gartenhumus, RICOTER Erdaufbereitung AG, Aarberg, Switzerland), to each plot to ensure the plants established well. Mesh fabric netting around each plot minimized the possibility of cross-pollination between the same species from different experimental plots. We collected seeds from seven monoculture plots, one 4-species mixture plot and six 8-species mixture plots in the experimental garden. We did not include seeds from 2-species mixtures as we expected that the community diversity selection pressure may not be different enough from monocultures. Seeds from different mother plants were pooled together and cleaned manually for three species and mechanically for two species (*P. lanceolata* and *P. vulgaris*). The dry seeds were stored for 5 months at 5° C for cold stratification.

### 2.4 Experimental set up

All seeds were germinated in germination soil (“Anzuchterde”, Ökohum, Herbertingen, Germany) under constant conditions in the glasshouse without additional light in December 2014. From 25 February to 13 March 2015, seedlings were planted in monocultures of four individuals or 2-species mixtures of two plus two individuals into pots (two liters) filled with neutral agricultural soil (50% agricultural sugar beet soil, 25% perlite, 25% sand; Ricoter AG, Aarberg, Switzerland). At the beginning of the experiment the studied plants were infested by fungus gnats (*Bradysia* spp.), which caused seedling mortality during the experiment. Seedlings that died in the first two weeks were replaced with seedlings of the same species and age.

We planted species assemblages in six blocks (replicates); each block included the full experimental design. Species pairs were chosen according to seedling availability. The full design (every possible species combination) was intended but could not be realized due to seedling mortality and low germination rates for some species (e.g. *G. mollugo*). Within each block, pots were placed on three different tables in the glasshouse in a randomized fashion without reference to selection history or assembly treatment. During the timeframe of the experiment we did not move pots but noted their position in the glasshouse. Single pots always contained four plants of the same selection history. Each selection history × species assembly combination was replicated five to six times depending on plant availability. We planted 30 monoculture and 42 mixture assemblages with mixture selection history, 30 monoculture and 60 mixture assemblages with monoculture selection history and 24 monoculture and 35 mixture assemblages with naïve selection history, a total of 221 pots and 884 plants (Appendix S2 for monoculture identities and species combinations).

During the experiment, the plants were initially kept at day temperatures of 17-20°C and night temperatures of 13-17°C without supplemental light. To compensate for overheating in summer, an adiabatic cooling system (Airwatech; Bern, Switzerland) was used to keep inside temperatures constant with outside air temperatures. The plants were not fertilized. Due to an infestation of white flies (*Trialeurodes vaporariorum*, Westwood 1856) and spider mites (*Tetranychidae* spp., Donnadieu 1875), we applied the insecticide SanoPlant Neem (1% Azadirachtin A (10 g/l); Maag AG) three times. The fungicide Fenicur (*Oleum foeniculi*, Andermatt Biocontrol) against powdery mildew (*Podosphaera* spp.) was applied twice. Plant height, leaf thickness, specific leaf area (SLA) and individual aboveground biomass were measured after twelve weeks of the experiment from 18 May to 4 June 2015. Leaf thickness was measured for three representative leaves using a thickness gauge. Specific leaf area (SLA) of up to 20 representative leaves (depending on the leaf size of the species) of each species in a pot was measured by scanning fresh leaves with a Li-3100 Area Meter (Li-cor Inc., Lincoln, Nebraska, USA) immediately after harvest and determining the mass of the same leaves after drying. Plant height and individual aboveground biomass were measured a second time after 24 weeks from 18-25 August 2015 at the end of the experiment. All four individuals in a pot were sampled. Research assistants, who were not informed of the specific experimental treatments, assisted in the regular measurements and harvesting of plants at the end of the experiment.

### 2.5 Data analysis

We calculated pot-wise aboveground community biomass (plant community production) as the sum of the biomass of the four individual plants. Because the first measure assessed growth and the second regrowth, the harvests were analyzed separately. Relative between-species differences (RDs, absolute difference between two species divided by the mean of the two) in plant height (first and second harvest), leaf thickness (first harvest) and SLA (first harvest) were calculated for mixture assemblages. Relative differences within species were calculated for both mixture and monoculture assemblages taking the relative difference between two individuals of the same species per pot. SLA outliers (> 99% percentile) were replaced with a maximum value (the 99% percentile, n = 6). Furthermore, we calculated community-weighted means (CWMs) and pot standard deviation (SDs) for the same traits (R package “FD", Laliberté and Legendre 2010; Laliberté et al., 2014). Pots with dead plant individuals were excluded from the calculation of community-weighted means, but were included for the other data analyses. Net biodiversity effects (NE) were calculated by comparing the 2-species mixtures with the average monoculture and partitioned according to Loreau and Hector (2001) into complementary (CE) and sampling (selection) effects (SE). This partitioning approach allows assessing how CE and SE contribute to the NE (Loreau and Hector 2001). To avoid confusion with the term selection used for the selection-history treatment, we here use the term “sampling effect” for the SE (see Zuppinger-Dingley et al., 2014). Additive partitioning calculations were based on the difference between the observed yield of each species in the mixture and the monoculture yield for that species and selection history averaged across blocks. Absolute values of CE and SE were square root-transformed and the original signs put back on the transformed values for analysis (Loreau and Hector 2001). Differences in these measures between mixtures assembled from plants with monoculture selection history and mixtures assembled from plants with mixture selection history would suggest differential evolution of trait-based niches between species as a potential mechanism underlying biodiversity effects.

All statistical analyses were done in R (Version 3.2.3, R Core team 2016). Mixed-model analysis was done using the R-package asreml (VSNI international, 2016) and results were assembled in ANOVA tables. Selection-history treatment (naïve, monoculture, mixture), assembly treatment (monoculture vs. 2-species mixture assemblages), species identity of monoculture assemblages and of mixture assemblages (in short “species assembly”) and interactions of these were fixed-effects terms in the model. Table (including blocks) was a random-effects term in the model. CWMs, RDs, within-species differences and SDs of plant height, SLA and leaf thickness were added as covariates to determine the influence of these covariates on community biomass and biodiversity effects.

## 3. Results

### 3.1 Plant selection history and community productivity

Assemblages consisting of plants with naïve selection history produced the lowest community biomass at both harvests (Fig. 1; Table 1). At the second harvest, such higher productivity of selected plants was stronger in 2-species mixtures than in monocultures (Fig. 1; Table 1). At the second harvest, mixture-type plant assemblages outperformed monoculture-type plant assemblages and this effect was marginally more pronounced in the 2-species mixtures (Fig. 1; Table 1).

**Table 1.** Results of mixed-effects ANOVA for the aboveground biomass of the test assemblages at first harvest after 12 weeks of growth **(a)** and at second harvest after 24 weeks of growth **(b).**

| Source of variation |  | nDf | a) Harvest 1 |  |  |  | b) Harvest 2 |  |  |
| --- | --- | --- | --- | --- | --- | --- | --- | --- | --- |
|  |  |  | dDF | *F* | *P* |  | dDF | *F* | *P* |
| Species assembly: |  |  |  |  |  |  |  |  |  |
|  | Monoculture vs. mixture | 1 | 173.3 | 29.09 | < 0.001 |  | 174 | 10.78 | 0.001 |
|  | Monoculture identity or species combination of mixture | 13 | 171.2 | 16.53 | < 0.001 |  | 171.8 | 15.47 | < 0.001 |
| Selection history: |  |  |  |  |  |  |  |  |  |
|  | Naïve vs. mono or mix types | 1 | 173 | 16.63 | < 0.001 |  | 173.7 | 42.72 | < 0.001 |
|  | Mono vs. mix types | 1 | 169.6 | 1.78 | 0.184 |  | 170.1 | 5.71 | 0.018 |
| Assembly × history: |  |  |  |  |  |  |  |  |  |
|  | Monoculture vs. mixture × naïve vs. mono or mix types | 1 | 168.4 | 1.72 | 0.191 |  | 168.8 | 8.56 | 0.004 |
|  | Monoculture vs. mixture × Mono or mix types | 1 | 172.2 | 1.69 | 0.195 |  | 172.9 | 3.52 | 0.062 |
| Species assembly × history: |  |  |  |  |  |  |  |  |  |
|  | Species assembly × naïve vs. mono or mix types | 8 | 171.7 | 5.35 | < 0.001 |  | 172.3 | 2.15 | 0.033 |
|  | Species assembly × mono types vs. mix types | 10 | 172.3 | 2.91 | 0.002 |  | 172.9 | 1.23 | 0.275 |

nDF = numerator degrees of freedom, dDF = denominator degrees of freedom, *F* = variance ratio, *P* = probability of type-I error. Variance components (Var) and associated standard errors (SE) for the random effects are provided together with the number of replicates.

**Fig. 1.**
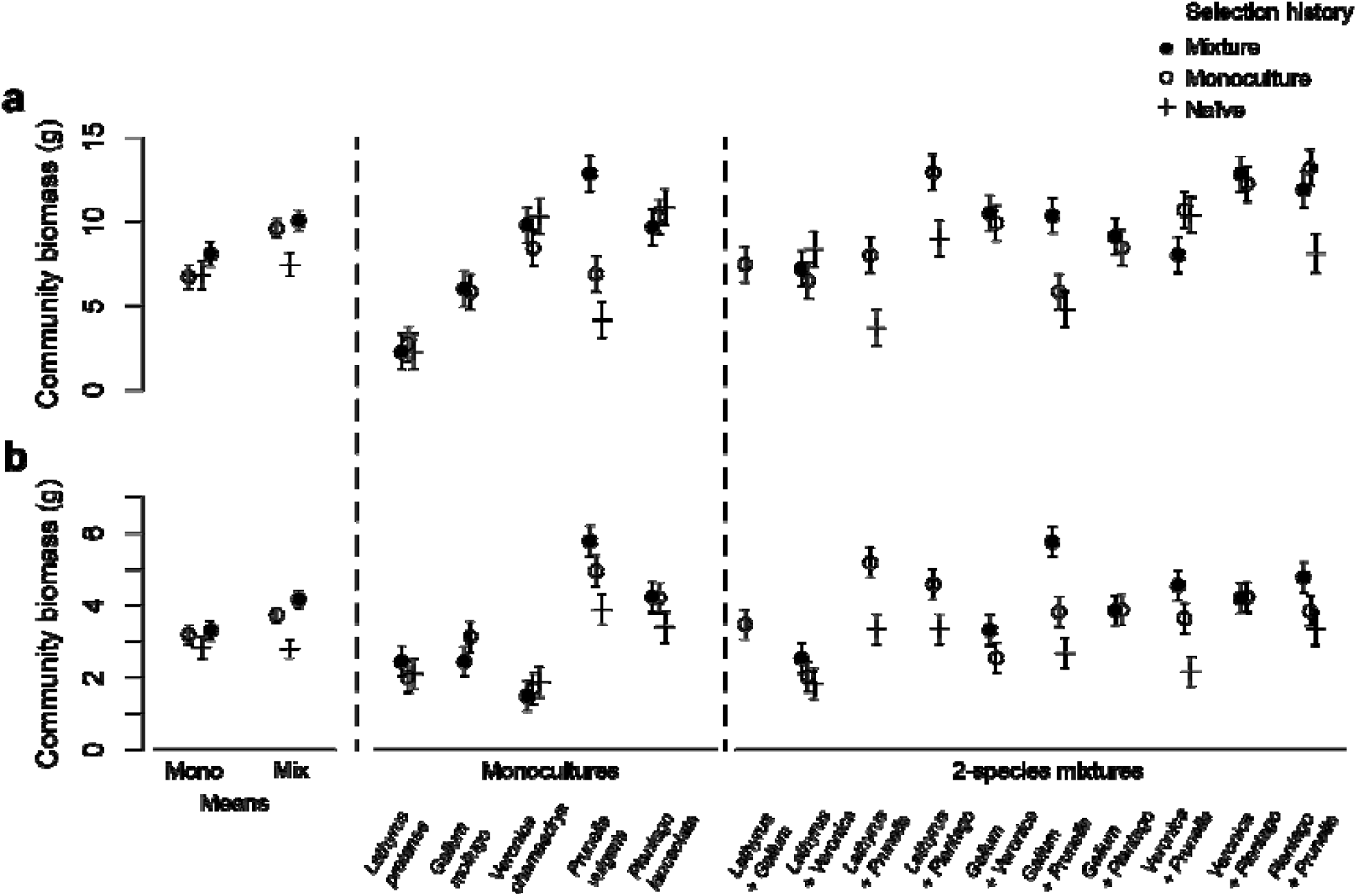
Mean community biomass for monocultures and 2-species mixtures. Shown are means and standard errors from a linear mixed-effects model with selection history, species combination and the interaction between selection history and species assembly as fixed-effects terms and table (including the block) as random-effects term. **a**, first harvest. **b**, second harvest.

Species identity in monoculture or mixture assemblages strongly influenced community productivity and, especially at the first harvest, the interaction terms with selection history were significant (Table 1). For example, at the first harvest, mixture-type plants performed better than monoculture-type plants in newly assembled monocultures of *P. vulgaris* and in mixtures of *G. mollugo* and *P. vulgaris* (Fig. 1a). However, in the two mixtures with the small herbs *V. chamaedrys* and *P. vulgaris* and *P. lanceolata* and *P. vulgaris*, monoculture-type plants performed better than mixture-type plants (see Fig. 1a).

### 3.2 Plant selection history and biodiversity effects

Overall, biodiversity effects were positive at both harvests (Fig. 2, Table 2). Communities of naïve plants at the first harvest showed larger SEs and at the second harvest showed smaller NEs and CEs than communities of selected plants.

**Fig. 2.**
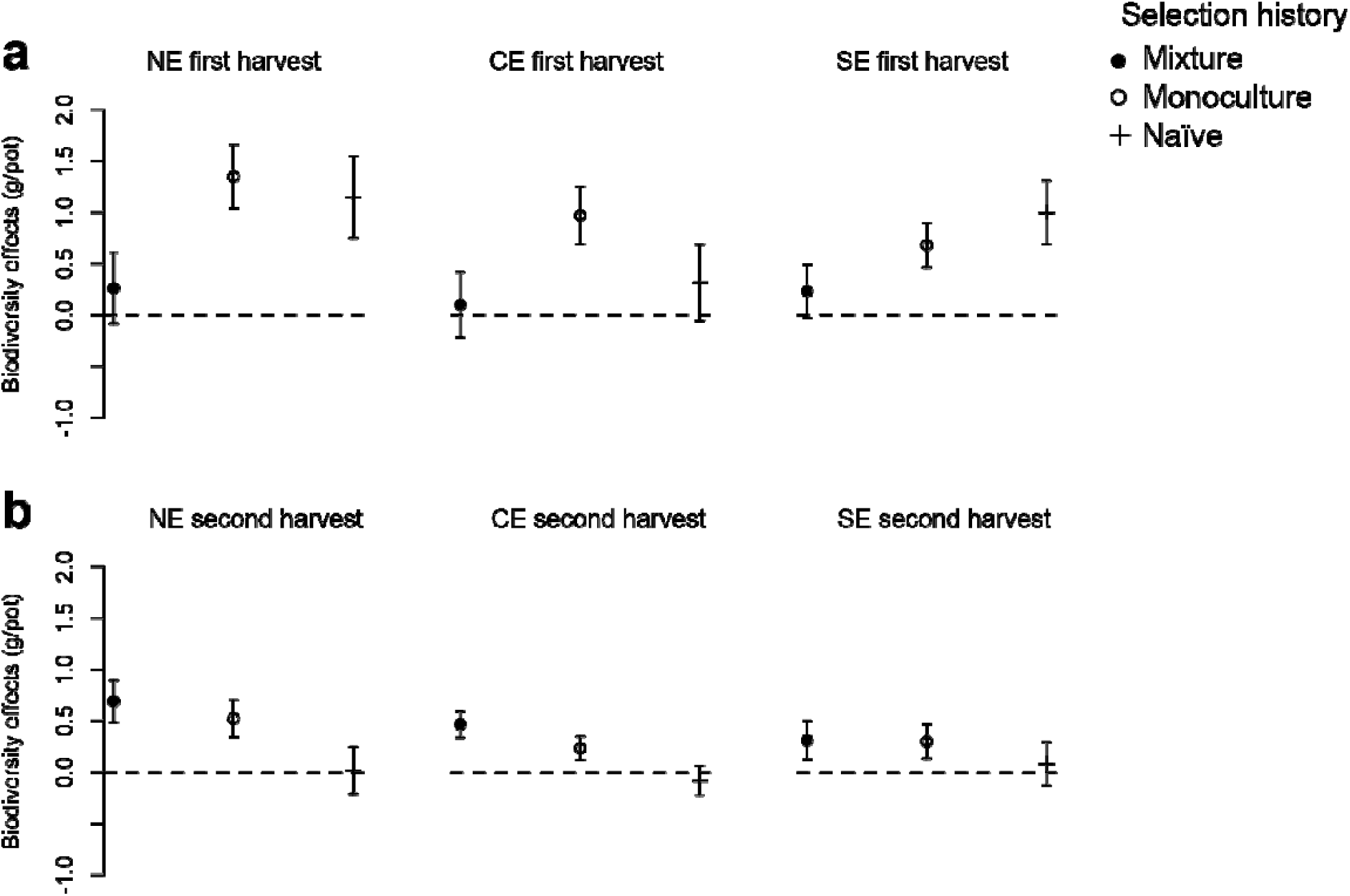
We assessed biodiversity effects for both biomass harvests by additive partitioning of the net effect (**a,** NE) into complementarity effect (**b,** CE) and sampling effect (**c,** SE) for plants with different selection histories (naïve, monoculture, mixture). Shown are means and standard errors from a linear mixed-effects model, with selection history as fixed-effects term and species assembly, the interaction between selection history and species assembly and table (including block) as random-effects terms.

**Table 2.** Results of mixed-effects ANOVA for biodiversity effects of the test assemblages at the first harvest after 12 weeks of growth **(a)** and at the second harvest after 24 weeks of growth **(b)**

| Source of variation | nDf | a) NE Harvest 1 |  |  | b) NE Harvest 2 |  |  |
| --- | --- | --- | --- | --- | --- | --- | --- |
|  |  | dDF | *F* | *P* | dDF | *F* | *P* |
| Overall mean | 1 | 15.9 | 26.67 | < 0.001 | 15.1 | 14.35 | 0.002 |
| Naïve vs. mono or mix types | 1 | 95.3 | 0.806 | 0.372 | 96.1 | 11.54 | < 0.001 |
| Mono types vs. mix types | 1 | 93.9 | 21.01 | < 0.001 | 93.6 | 0.026 | 0.872 |
| Species assembly | 9 | 96.7 | 2.646 | 0.009 | 97.7 | 4.837 | < 0.001 |
| Species assembly × naïve vs. mono or mix types | 4 | 97.5 | 4.459 | 0.002 | 98.9 | 1.463 | 0.219 |
| Species assembly × mono types vs. mix types | 6 | 98 | 4.095 | 0.001 | 99 | 1.518 | 0.18 |

| Source of variation | nDf | a) CE harvest 1 |  |  | b) CE Harvest 2 |  |  |
| --- | --- | --- | --- | --- | --- | --- | --- |
|  |  | dDF | *F* | *P* | dDF | *F* | *P* |
| Overall mean | 1 | 15.8 | 8.214 | 0.011 | 14.5 | 4.108 | 0.061 |
| Naïve vs. mono or mix types | 1 | 95.9 | 1.427 | 0.235 | 96.6 | 5.668 | 0.019 |
| Mono types vs. mix types | 1 | 94.4 | 14.2 | < 0.001 | 93.8 | 1.524 | 0.22 |
| Species assembly | 9 | 97.4 | 2.534 | 0.012 | 98.4 | 1.121 | 0.356 |
| Species assembly × naïve vs. mono or mix types | 4 | 98.3 | 1.835 | 0.128 | 99.7 | 0.584 | 0.675 |
| Species assembly × mono types vs. mix types | 6 | 98.8 | 2.53 | 0.025 | 99.8 | 0.468 | 0.831 |

| Source of variation | nDf | a) SE harvest 1 |  |  | b) SE Harvest 2 |  |  |
| --- | --- | --- | --- | --- | --- | --- | --- |
|  |  | dDF | *F* | *P* | dDF | *F* | *P* |
| Overall mean | 1 | 14.2 | 97.07 | < 0.001 | 15.1 | 11.66 | 0.004 |
| Naïve vs. mono or mix types | 1 | 104.2 | 12.66 | 0.001 | 98.8 | 2.224 | 0.139 |
| Mono types vs. mix types | 1 | 101.2 | 10.28 | 0.002 | 95.7 | 2.37 | 0.127 |
| Species assembly | 9 | 105.5 | 5.793 | < 0.001 | 100.8 | 11.53 | < 0.001 |
| Species assembly × naïve vs. mono or mix types | 4 | 105.9 | 10.08 | < 0.001 | 102 | 3.517 | 0.01 |
| Species assembly × mono types vs. mix types | 6 | 105.9 | 2.865 | 0.013 | 101.9 | 2.541 | 0.025 |

nDF = numerator degrees of freedom, dDF = denominator degrees of freedom, *F* = variance ratio, *P* = probability of type-I error. Variance components (Var) and associated standard errors (SE) for the random effects are provided together with the number of replicates.

At the first harvest, NEs, CEs and SEs were also larger for communities assembled from monoculture-type plants than for communities assembled from mixture-type plants (Fig. 2a, Table 2a). In contrast, at the second harvest NEs, CEs and SEs were non-significantly larger for communities assembled from mixture-type plants for most species assemblages (Fig. 2b; Table 2b). As with the results obtained for community productivity, the influence of selection history on biodiversity effects also depended on the specific species combination in mixture assemblages (Table 2). NEs were larger for mixture-type plants for the combinations of *G. mollugo* with either *P. vulgaris* or *P. lanceolata* at the first harvest (Fig. A2 in Supplementary material; Table 2). At the second harvest, NEs and CEs were generally more similar between selection histories across different combinations and variation between the specific community compositions was mainly due to different SEs. The much larger NE for mixture-type plants in the combination *G. mollugo* + *P. vulgaris* was an exception (Fig. A2). For both harvests communities which included the legume *L. pratensis* or the small herb *P. lanceolata* showed positive biodiversity effects (Fig. A2). Four species combinations shifted between harvests from stronger CEs for monoculture-type plants to stronger biodiversity effects for mixture-type plants (Fig. A2). The *G. mollugo + P. vulgaris* combination showed a consistently larger CE for mixture-type plants. At the second harvest the different species combinations varied strongly in SEs, but not in CEs (Table 2). SEs were often larger for mixture- than for monoculture-type plants (Fig. 2).

### 3.3 Plant selection history and within- and between-species trait variance

Whereas interspecific differences in plant height were marginally larger in mixture-type plants, interspecific differences in leaf thickness were larger in monoculture-type plants at the first harvest (Fig. 3). Intraspecific differences in SLA were larger for monoculture-type plants. Furthermore, pot-level SDs in monoculture or mixture assemblages were non-significantly larger for assemblages with monoculture- than with mixture-type plants (left two columns in Fig. 3).

**Fig. 3.**
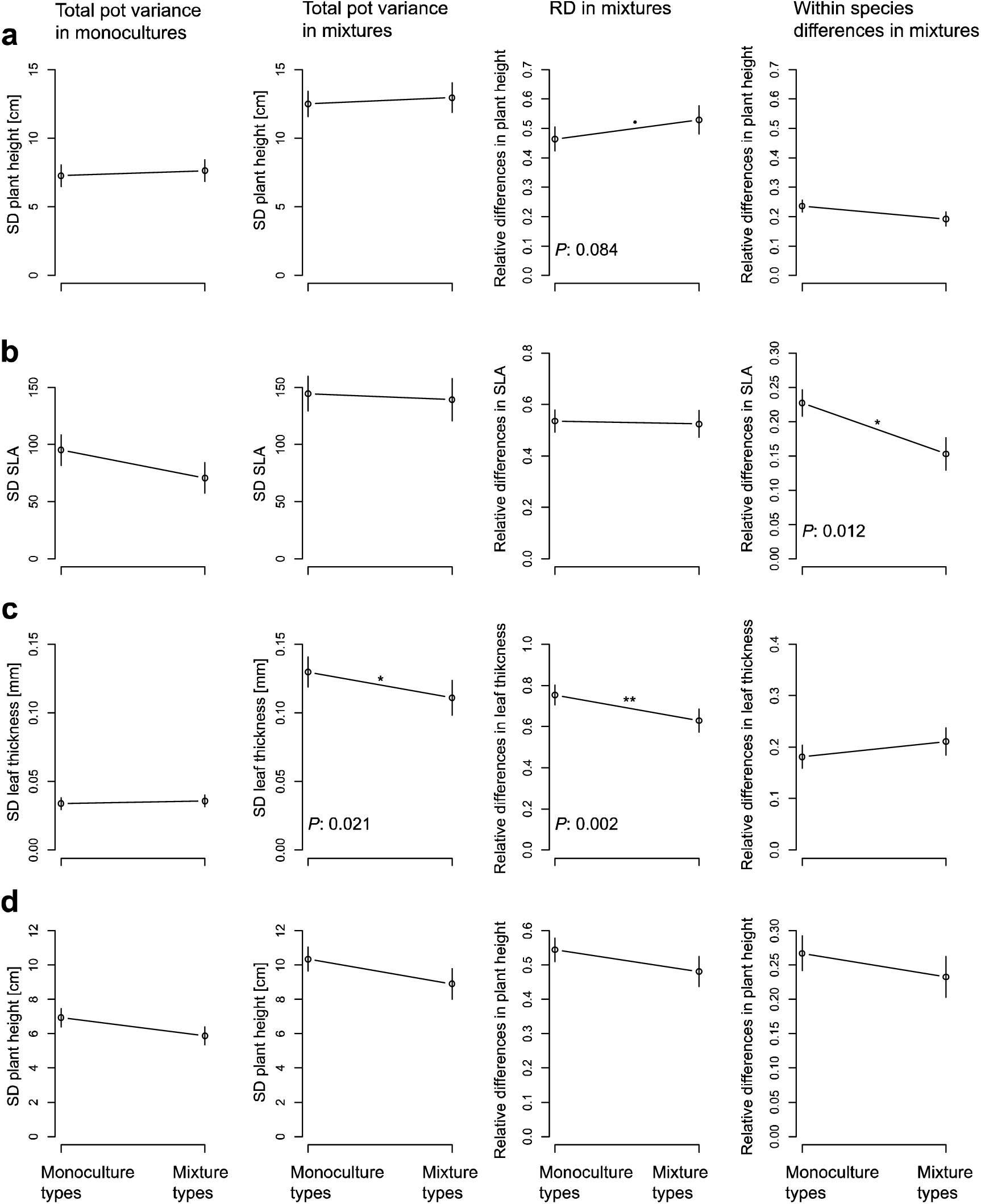
Trait variance in monoculture and mixture assemblages in response to selection history (monoculture- vs. mixture-type plants). **a)** plant height at first harvest, **b)** SLA at the first harvest, **c)** leaf thickness at the first harvest, **d)** plant height at second harvest. Shown are means and standard errors from a mixed-effects model with selection history, species assembly and the two-way interaction of these as fixed-effects terms and table (including block) as random term. Significant and marginally significant *P*-values are indicated in the respective plot.

### 3.4 Relationship between biodiversity effects and trait variation

At the first harvest, the NE was negatively correlated with the RD of plant height but positively correlated with the RD of leaf thickness (Fig. 4). Selection history had a significant effect on the relationship between SE and the RD of plant height but no or only marginal effects on all other relationships. SEs were more negatively correlated with the RDs of plant height for mixture- than for monoculture-type or naïve plants. In contrast, the RDs of leaf thickness were positively correlated with NEs and CEs for both monoculture- and mixture-type plants, but not for naïve plants (Fig. 4c). At the second harvest, NEs and SEs were significantly negatively correlated with the RD of plant height (Fig. 5). CEs were not influenced by interspecific variation in plant height.

**Fig. 4.**
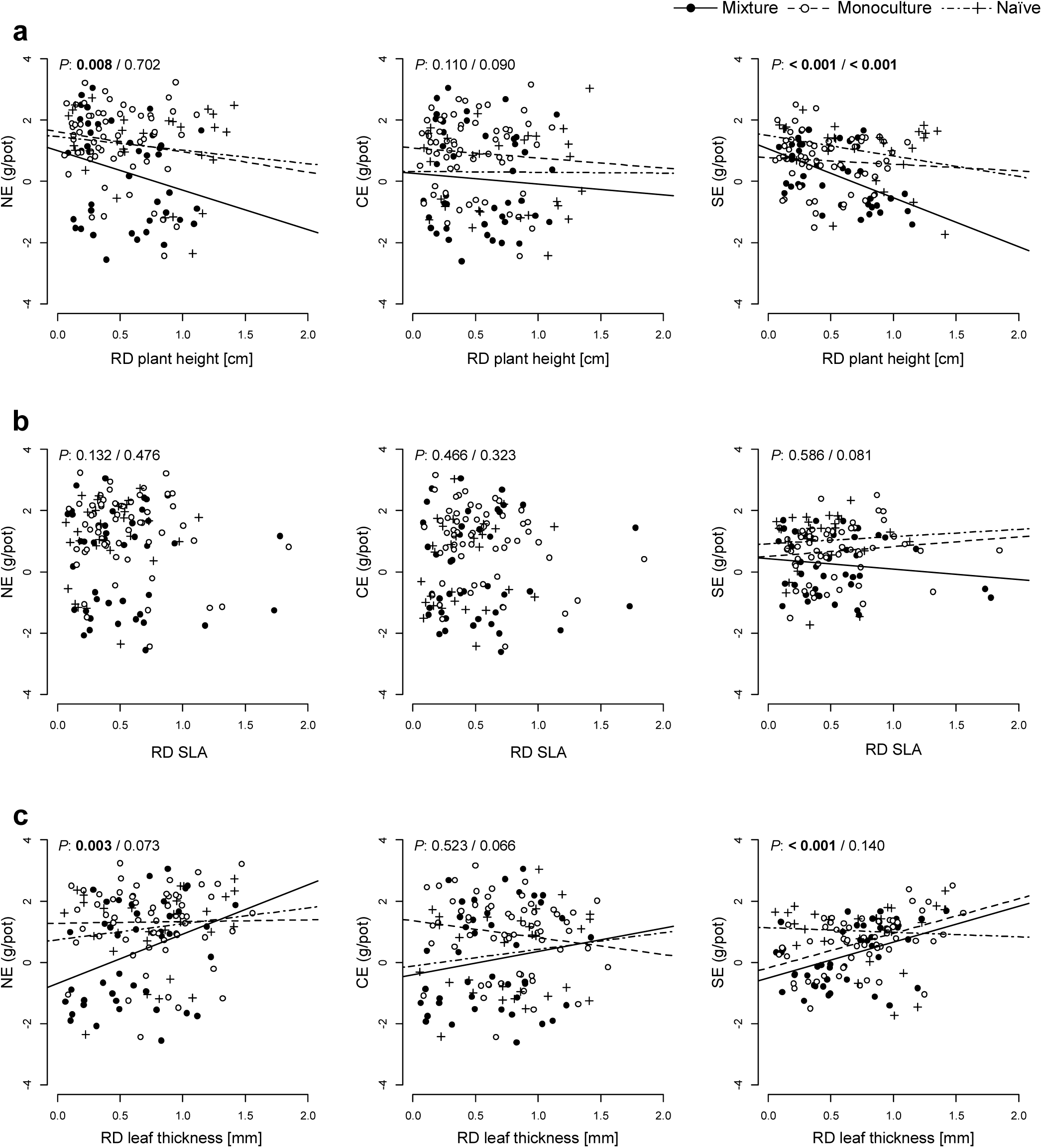
Biodiversity effects at the first harvest in response to relative differences between species (RDs) for three traits: **a,** plant height (in cm), **b,** specific leaf area (SLA) and **c,** leaf thickness (in mm). Indicated *P*-values refer to ANOVA results for fixed-effects terms from a mixed-effects model with RD, species assembly, selection history and interactions of these as fixed-effects terms and table (including block) as random-effects term: RD / interaction RD × selection history (naïve plants vs. mixture types vs. monoculture types). Regression lines are plotted in cases for which at least one *P*-value was significant. Left column: NE, middle column: CE, right column: SE.

**Fig. 5.**
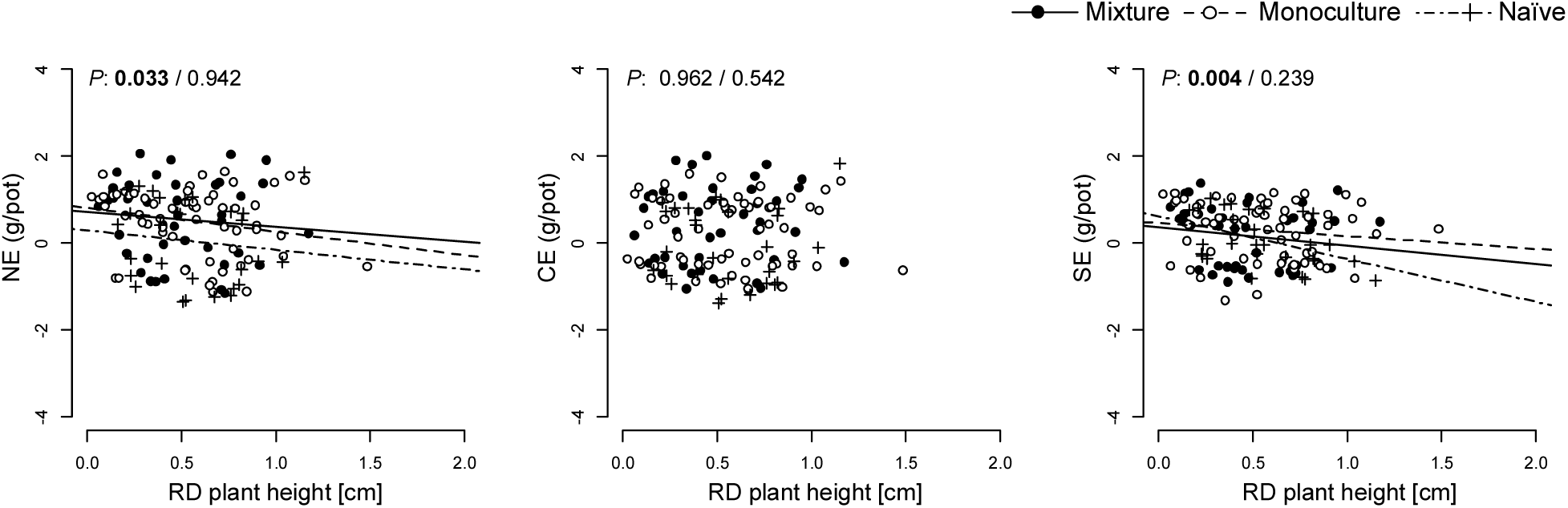
Biodiversity effects at the second harvest in response to relative differences between species for plant height (in cm). Indicated *P*-values refer to ANOVA results for fixed-effects terms from a mixed-effects model with RD, species assembly, selection history and interactions of these as fixed-effects terms and table (including block) as random-effects term: RD / interaction RD × selection history (naïve plants vs. mixture types vs. monoculture types). Regression lines are plotted in cases for which at least one *P*-value was significant. Left column: NE, middle column: CE, right column: SE.

### 3.5 Relationship between biodiversity effects and trait means

Whereas the CE was negatively correlated with the CWM of SLA (Fig. 6b), the SE was positively correlated with the CWM of SLA (Fig. 6b, right panel). Consequently, the NE, driven by the CE, decreased with increasing SLA. Leaf thickness had a marginally significant effect on SE, but the directionality depended on selection history. Plant height did not have a significant effect on the biodiversity effects at the first harvest. However, the interaction between trait means and selection history was significant for the relationship between the CWM of plant height and the SE at the first harvest. Selection history was not significant for the relationship between biodiversity effects and CWMs for the other two traits. At the second harvest, CWM of plant height had a significantly positive effect on NE, CE and SE (Fig. 7), the biodiversity effects were therefore stronger for taller plants. However, in contrast to the first harvest, at the second harvest no effect of selection history on the relationship between the CWM of plant height and the SE was observed (Fig. 7).

**FIG. 6.**
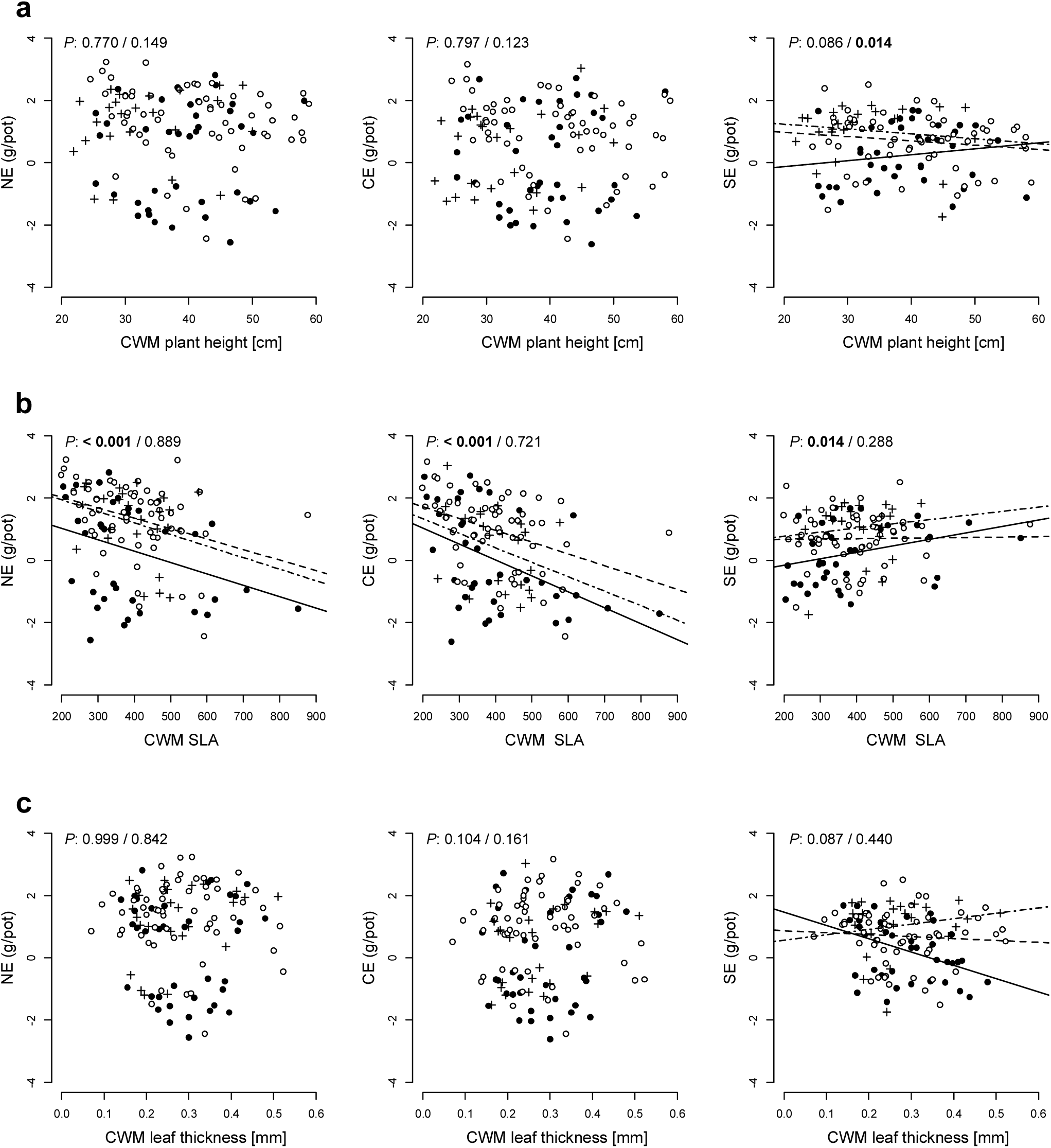
Biodiversity effects at the first harvest in response to the community-weighted mean (CWM) of three traits: **a,** plant height (in cm), **b,** specific leaf area (SLA) and **c,** leaf thickness (in mm). Indicated *P*-values refer to ANOVA results for fixed-effects terms from a mixed-effects model with CWM, species assembly, selection history and interactions of these as fixed-effects terms and table (including block) as random-effects term: CWM / interaction CWM × selection history (naïve plants vs. mixture types vs. monoculture types). Regression lines are plotted in cases for which at least one *P*-value was significant. Left column: NE, middle column: CE, right column: SE.

**Fig. 7.**
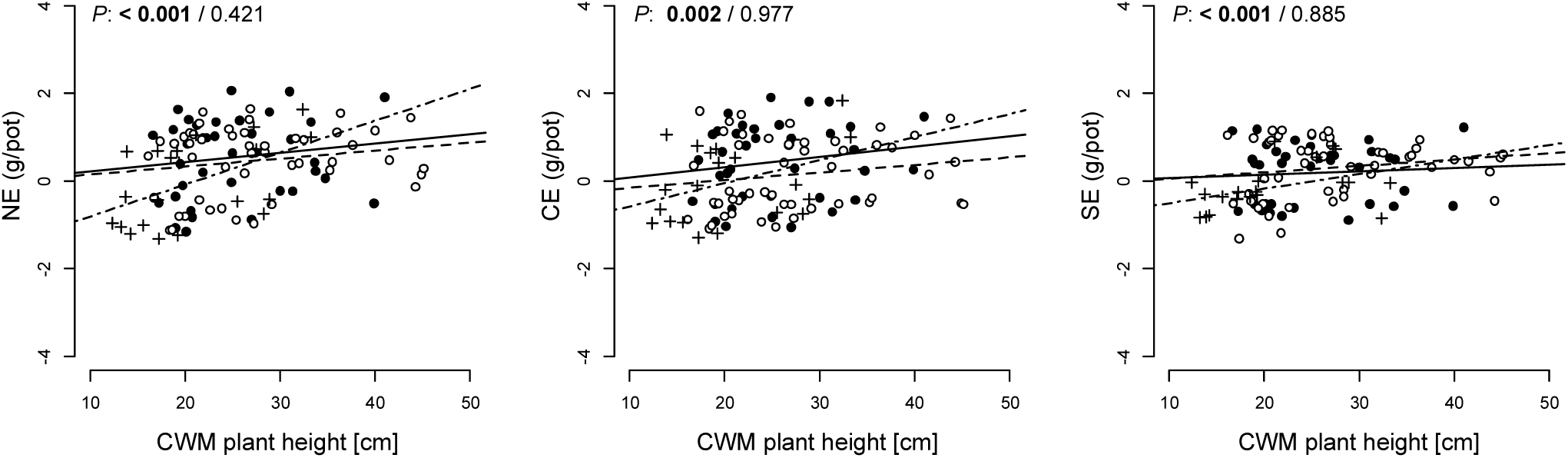
Biodiversity effects at the second harvest in response to the community-weighted mean (CWM) of plant height (in cm). Indicated *P*-values refer to ANOVA results for fixed-effects terms from a mixed-effects model with CWM, species assembly, selection history and interactions of these as fixed-effects terms and table (including block) as random-effects term: CWM / interaction CWM × selection history (naïve plants vs. mixture types vs. monoculture types). Regression lines are plotted in cases for which at least one *P*-value was significant. Left column: NE, middle column: CE, right column: SE.

## 4. Discussion

### 4.1 Influence of plant selection history on community productivity

Plant community productivity may be influenced by selection for increased niche differentiation in plants grown for eight years in mixtures (mixture-type plants) compared with plants grown in monocultures (monoculture-type plants, Zuppinger-Dingley et al., 2014). We hypothesized that 2-species mixtures comprised of mixture-type plants should have greater community productivity than the same mixtures comprised of monoculture-type plants. Conversely, we expected monocultures with monoculture-type plants to have greater community productivity than the same monocultures with mixture-type plants. For naïve plants, we expected intermediate community productivity in both monocultures and mixtures.

Our results provide mixed evidence for these hypotheses, in part depending on the particular species and species combinations. Plant assemblages consisting of naïve plants, without a selection history in the Jena Experiment, generally produced the lowest community biomass, especially in 2-species mixtures, in the pots in the glasshouse in Zurich. Evolutionary processes in the field plots likely led to the increased performance of selected plants, because these plants were grown for a longer time without re-sowing. In contrast, the naïve plants were re-sown every year in the commercial propagation cultures, thereby “resetting” any local adaptation with every generation.

Within the selected plants, mixture-type plants produced higher community biomass than monoculture-type plants in 2-species mixtures. But mixture-type plants also produced more biomass than monoculture-type plants when grown in monoculture, which reduced biodiversity effects. The generally lower performance of monoculture-type plants may have been due to selection for increased defense against pathogens that are known to accumulate in monocultures (Schnitzer et al., 2011). Increased resource allocation to defense mechanisms would result in reduced allocation to growth (Coley et al., 1985; Herms and Mattson, 1992). However, during the 24 weeks of growth in the experiment, species-specific pathogens may not have been present, removing the need for increased defense. Such species-specific pathogens might have needed more time to accumulate and to render increased pathogen resistance advantageous.

Selection-history effects varied strongly among species in monocultures and among species compositions in 2-species mixtures. These findings emphasized that such studies should include multiple species but at the same time sufficient replication in monoculture and their combinations in mixture. High replication can more easily be achieved in experiments with one focal species (e.g. Kleynhans et al., 2016; Rottstock et al., 2017), but extrapolating results from such experiments might under- or overestimate overall effects of selection on the response of plants to different biotic conditions. We used five focal species and observed strong differences regarding their selection response to community diversity.

### 4.2 Influence of plant selection history on biodiversity effects

Net biodiversity effects can be partitioned into CEs and SEs. When CEs drive over-yielding, most species contribute similarly to greater community productivity in mixtures, presumably due to niche differentiation among them. Conversely, SEs are large when few dominant species are driving positive diversity-productivity relationships, because they benefit from growing in mixtures (Loreau and Hector 2001).

Naïve plants exhibited weak biodiversity effects, confirming findings from a field experiment, where biodiversity effects were weaker for assemblages of naïve plants, especially at low diversity levels (van Moorsel et al., 2017). Such naïve plants, in contrast to plants with a common selection history, did not experience continued selection in field plots without re-sowing and did not previously experience interspecific competition. In contrast to our expectations, not only NEs and SEs but also CEs were larger for monoculture- than for mixture-type plant communities at the first harvest. However, the lower CEs of mixture-type communities could be attributed to a higher performance of mixture-type plants in monoculture assemblages and not to a lower performance in mixed assemblages. At second harvest, NEs, CEs and SEs were similar for mixture- and monoculture-type communities. Nevertheless, at least in four 2-species combinations – *L. pratensis* + *V. chamaedrys*, *G. mollugo* + *V. chamaedrys*, *V. chamaedrys* + *P. vulgaris* and *P. lanceolata* + *P. vulgaris* – the directionality changed, i.e. CEs at the second harvest were larger for mixture- than for monoculture-type communities. Over longer timespans, CEs often increase and SEs often decrease (Fargione et al., 2007; Isbell et al., 2009; Marquard et al., 2009; Montès et al., 2008; van Ruijven and Berendse, 2005). Possibly this would have occurred in our experiment if we had continued beyond the 24-weeks of the study.

### 4.3 Influence of plant selection history on trait variation

Because community-level trait variation can reflect niche differentiation (Roscher et al., 2015; Violle et al., 2012), we measured intra- and interspecific trait variation among individual plants in all assemblages. We expected larger interspecific trait variation for mixture-type plants undergoing possible selection for increased complementarity during twelve years in the experimental field plots. Conversely, we expected stronger within-species trait variation in monoculture-type plants with twelve years of strong intraspecific competition in the experimental field plots. However, monoculture-type plants tended to show higher intra and inter-specific trait variation (see Fig. 3). The relative extent of intraspecific trait variation may depend on species richness (Hulshof et al., 2013; Lamanna et al., 2014; Le Bagousse-Pinguet et al., 2014; Siefert et al., 2015) and in monocultures a large intraspecific variation is advantageous for a more efficient use of resources. Thus, the trend for increased trait variation in monoculture-type plants is consistent with potential selection for intraspecific niche widening by within-species character displacement during prolonged growth in monocultures.

However, less interspecific trait variation in mixture- compared with monoculture-type plants was in accordance with the lower CEs for mixture-type plants. These findings contrast with an earlier study in which larger CEs were observed for mixture- than for monoculture type plants and where mixture-type plants showed increased interspecific trait variation (Zuppinger et al., 2014). This earlier study included more species that were functionally different from each other, namely grasses, legumes, small herbs and tall herbs, which could potentially explain the contrasting results. More similar species, such as those used in the present study, may not be able to further increase trait differences in such a short time frame (Allan et al., 2013). Such species may have evolved “parallel” character displacement in response to species of the other functional groups also present in the mixtures in which they were selected in the Jena Experiment.

### 4.4 Influence of trait variation and community-weighted means on biodiversity effects

Selection for niche differentiation (Zuppinger-Dingley et al., 2014) could explain the increase of biodiversity effects over time in field experiments (Cardinale et al., 2007, Reich et al., 2012). Not all trait variation, however, corresponds to niche differentiation (Turcotte and Levine 2016). In particular, traits related to light availability may behave differently because of the asymmetric nature of competition for light, i.e. being tall is generally better than being small. Thus, variation in plant height could be expected to decrease when species are grown in mixtures rather than monocultures (Roscher et al., 2015; Vermeulen et al., 2008). In the present study, the relationship between functional traits in 2-species mixtures and biodiversity effects did not consistently differ between plants selected in monocultures vs. mixtures and this was consistent with the absence of increased CEs and between-species trait variations in mixture-type plants. Nevertheless, we did test for such relationships – independent of selection treatments – between functional traits and biodiversity effects in our 2-species mixtures. Specifically, we tested whether relative trait differences (RD) were positively correlated with CEs and community-weighted trait means (CWMs) were positively correlated with SEs.

RDs in plant height were negatively rather than positively correlated with CEs and consequently NEs (see Fig. 4a, 5). This discrepancy between our expectation and observation suggests that RDs in plant height may reflect competitive hierarchies rather than complementary of plants with respect to light use, as discussed above with regard to the asymmetry of light competition. At the second harvest, CWMs of plant height had a positive impact on all biodiversity effects (Fig. 7), i.e. not only as has been previously observed on SEs (Cadotte 2017; Roscher et al., 2015; Vermeulen et al., 2008).

Functional diversity in SLA within a community should increase complementary light use (Roscher et al., 2011). Leaf thickness is inherently related to SLA (White and Montes-R, 2005) and might act similarly to SLA. Here, RDs in leaf thickness, but not RDs in SLA, were positively correlated with all biodiversity effects, especially for mixture-type plants. Trait plasticity in leaf thickness was therefore advantageous for species growing in mixtures. However, SEs increased as much as CEs, contrary to our expectation that positive correlations between trait differences may mainly involve CEs. Furthermore, CWMs of SLA had a positive effect on SEs, but also a negative effect on CEs, adding up to a negative effect on NEs, suggesting that a smaller leaf area per unit mass for species growing in mixtures has a positive effect on productivity.

## 5. Conclusions

Here, we demonstrated that community diversity had the selective potential to alter species performances, which may in part explain the strengthening of biodiversity-ecosystem functioning relationship observed in field experiments (e.g. Reich et al., 2012). Selection in a biodiversity experiment increased community productivity in newly assembled test assemblages compared to assemblages composed of naïve plants without such selection history. Moreover, previous selection in mixtures increased community productivity in newly assembled mixtures and monocultures compared with previous selection in monocultures. These findings imply that co-evolutionary processes occurred in the 12-year selection period in the experimental plots of the biodiversity experiment and involving at least two sexual reproduction cycles. Such rapid evolutionary processes in grassland plant communities have implications for conservation strategies. Thus, it may not be sufficient to only conserve species in isolation but rather in communities or populations of species with co-evolved interactions.

## Acknowledgements

We are grateful to T. Zwimpfer, M. Furler, D. Trujillo and D. Topalovic for technical assistance and E. de Luca, N. Castro and M. Brezzi for help with data measurements. This study was supported by the Swiss National Science Foundation (grants number 147092 and 166457 to B. Schmid) and the University Research Priority Program Global Change and Biodiversity of the University of Zurich. The Jena Experiment is supported by the German Science Foundation (FOR 1451, SCHM 1628/5-2). S.J. van Moorsel, B. Schmid and D. Zuppinger-Dingley conceived the study, S.J. van Moorsel and T. Hahl carried out the experiment and S.J. van Moorsel, M.W. Schmid and B. Schmid analyzed the data. S.J. van Moorsel and B. Schmid wrote the manuscript with all other authors contributing to revisions.

## Supplementary data

Supplementary data associated with this article can be found in the online version.

## References

Aarssen, L.W., 1983. Ecological Combining Ability and Competitive Combining Ability in Plants: Toward a General Evolutionary Theory of Coexistence in Systems of Competition. Am. Nat. 122, 707–731.

Allan, E., Jenkins, T., Fergus, A.J., Roscher, C., Fischer, M., Petermann, J., Weisser, W.W., Schmid, B., 2013. Experimental plant communities develop phylogenetically overdispersed abundance distributions during assembly. Ecology 94, 465–477.

Allan, E., Weisser, W., Weigelt, A., Roscher, C., Fischer, M., Hillebrand, H., 2011. More diverse plant communities have higher functioning over time due to turnover in complementary dominant species. Proc. Natl. Acad. Sci. 108, 17034–17039.

Anderson, J.T., Willis, J.H., Mitchell-Olds, T., 2011. Evolutionary genetics of plant adaptation. Trends Genet. 27, 258–266.

Bossdorf, O., Richards, C.L., Pigliucci, M., 2008. Epigenetics for ecologists. Ecol. Lett. 11, 106–115.

Cadotte, M.W., 2017. Functional traits explain ecosystem function through opposing mechanisms. Ecol. Lett. 20, 989–996.

Cadotte, M.W., Cavender-Bares, J., Tilman, D., Oakley, T.H., 2009. Using Phylogenetic, Functional and Trait Diversity to Understand Patterns of Plant Community Productivity. PLoS ONE 4, e5695.

Cardinale, B.J., Duffy, J.E., Gonzalez, A., Hooper, D.U., Perrings, C., Venail, P., Narwani, A., Mace, G.M., Tilman, D., Wardle, D.A., Kinzig, A.P., Daily, G.C., Loreau, M., Grace, J.B., Larigauderie, A., Srivastava, D.S., Naeem, S., 2012. Biodiversity loss and its impact on humanity. Nature 486, 59–67.

Cardinale, B.J., Wright, J.P., Cadotte, M.W., Carroll, I.T., Hector, A., Srivastava, D.S., Loreau, M., Weis, J.J., 2007. Impacts of plant diversity on biomass production increase through time because of species complementarity. Proc. Natl. Acad. Sci. 104, 18123–18128.

Coley, P.D., Bryant, John P., Chapin, F. Stuart, 1985. Resource availability and plant antiherbivore defense. Science 230, 895–899.

Fakheran, S., Paul-Victor, C., Heichinger, C., Schmid, B., Grossniklaus, U., Turnbull, L.A., 2010. Adaptation and extinction in experimentally fragmented landscapes. Proc. Natl. Acad. Sci. 107, 19120–19125.

Fargione, J., Tilman, D., Dybzinski, R., Lambers, J.H.R., Clark, C., Harpole, W.S., Knops, J.M.., Reich, P.B., Loreau, M., 2007. From selection to complementarity: shifts in the causes of biodiversity-productivity relationships in a long-term biodiversity experiment. Proc. R. Soc. B Biol. Sci. 274, 871–876.

Flynn, D.F., Mirotchnick, N., Jain, M., Palmer, M.I., Naeem, S., 2011. Functional and phylogenetic diversity as predictors of biodiversity-ecosystem-function relationships. Ecology 92, 1573–1581.

Fornara, D.A., Tilman, D., 2008. Plant functional composition influences rates of soil carbon and nitrogen accumulation. J. Ecol. 96, 314–322.

Harper, J.L., 1977. Population biology of plants. London: Academic Press.

Hart, S.P., Schreiber, S.J., Levine, J.M., 2016. How variation between individuals affects species coexistence. Ecol. Lett. 19, 825–838.

Herms, D.A., Mattson, W.J., 1992. The Dilemma of Plants: To Grow or Defend. Q. Rev. Biol. 67, 283–335.

Hulshof, C.M., Violle, C., Spasojevic, M.J., McGill, B., Damschen, E., Harrison, S., Enquist, B.J., 2013. Intra-specific and inter-specific variation in specific leaf area reveal the importance of abiotic and biotic drivers of species diversity across elevation and latitude. J. Veg. Sci. 24, 921–931.

Isbell, F., Calcagno, V., Hector, A., Connolly, J., Harpole, W.S., Reich, P.B., Scherer-Lorenzen, M., Schmid, B., Tilman, D., van Ruijven, J., Weigelt, A., Wilsey, B.J., Zavaleta, E.S., Loreau, M., 2011. High plant diversity is needed to maintain ecosystem services. Nature 477, 199–202.

Isbell, F.I., Polley, H.W., Wilsey, B.J., 2009. Species interaction mechanisms maintain grassland plant species diversity. Ecology 90, 1821–1830.

Joshi, J., Schmid, B., Caldeira, M.C., Dimitrakopoulos, P.G., Good, J., Harris, R., Hector, A., Huss-Danell, K., Jumpponen, A., Minns, A., others, 2001. Local adaptation enhances performance of common plant species. Ecol. Lett. 4, 536–544.

Kleynhans, E.J., Otto, S.P., Reich, P.B., Vellend, M., 2016. Adaptation to elevated CO2 in different biodiversity contexts. Nat. Commun. 7, 12358.

Kraft, N.J.B., Godoy, O., Levine, J.M., 2015. Plant functional traits and the multidimensional nature of species coexistence. Proc. Natl. Acad. Sci. 112, 797–802.

Laliberté, E., Legendre, P., 2010. A distance-based framework for measuring functional diversity from multiple traits. Ecology 91, 299–305.

Laliberté, E., Legendre, P., Shipley, B., 2014. FD: measuring functional diversity from multiple traits, and other tools for functional ecology. R package version 1.0-12.

Lamanna, C., Blonder, B., Violle, C., Kraft, N.J.B., Sandel, B., imova, I., Donoghue, J.C., Svenning, J.-C., McGill, B.J., Boyle, B., Buzzard, V., Dolins, S., Jorgensen, P.M., Marcuse-Kubitza, A., Morueta-Holme, N., Peet, R.K., Piel, W.H., Regetz, J., Schildhauer, M., Spencer, N., Thiers, B., Wiser, S.K., Enquist, B.J., 2014. Functional trait space and the latitudinal diversity gradient. Proc. Natl. Acad. Sci. 111, 13745–13750.

Le Bagousse-Pinguet, Y., Xiao, S., Brooker, R.W., Gross, N., Liancourt, P., Straile, D., Michalet, R., 2014. Facilitation displaces hotspots of diversity and allows communities to persist in heavily stressed and disturbed environments. J. Veg. Sci. 25, 66–76.

Lipowsky, A., Schmid, B., Roscher, C., 2011. Selection for monoculture and mixture genotypes in a biodiversity experiment. Basic Appl. Ecol. 12, 360–371.

Loreau, M., Hector, A., 2001. Partitioning selection and complementarity in biodiversity experiments. Nature 72–76.

Marquard, E., Weigelt, A., Roscher, C., Gubsch, M., Lipowsky, A., Schmid, B., 2009. Positive biodiversity-productivity relationship due to increased plant density. J. Ecol. 97, 696–704.

Montès, N., Maestre, F.T., Ballini, C., Baldy, V., Gauquelin, T., Planquette, M., Greff, S., Dupouyet, S., Perret, J.-B., 2008. On the Relative Importance of the Effects of Selection and Complementarity as Drivers of Diversity-Productivity Relationships in Mediterranean Shrublands. Oikos 117, 1345–1350.

Mueller, K.E., Tilman, D., Fornara, D.A., Hobbie, S.E., 2013. Root depth distribution and the diversity-productivity relationship in a long-term grassland experiment. Ecology 94, 787–793.

Niklaus, P.A., Baruffol, M., He, J.-S., Ma, K., Schmid, B., 2017. Can niche plasticity promote biodiversity-productivity relationships through increased complementarity? Ecology.

Price, T.D., Qvarnstrom, A., Irwin, D.E., 2003. The role of phenotypic plasticity in driving genetic evolution. Proc. R. Soc. B Biol. Sci. 270, 1433–1440.

Reich, P.B., Tilman, D., Isbell, F., Mueller, K., Hobbie, S.E., Flynn, D.F.B., Eisenhauer, N., 2012. Impacts of biodiversity loss escalate through time as redundancy fades. Science 336, 589–592

Roscher, C., Kutsch, W.L., Schulze, E.-D., 2011. Light and nitrogen competition limit Lolium perenne in experimental grasslands of increasing plant diversity. Plant Biol. 13, 134–144.

Roscher, C., Schumacher, J., Baade, J., Wilcke, W., Gleixner, G., Weisser, W.W., Schmid, B., Schulze, E.-D., 2004. The role of biodiversity for element cycling and trophic interactions: an experimental approach in a grassland community. Basic Appl. Ecol. 5, 107–121.

Roscher, C., Schumacher, J., Gubsch, M., Lipowsky, A., Weigelt, A., Buchmann, N., Schmid, B., Schulze, E.-D., 2012. Using Plant Functional Traits to Explain Diversity-Productivity Relationships. PLoS ONE 7, e36760.

Roscher, C., Schumacher, J., Schmid, B., Schulze, E.-D., 2015. Contrasting effects of intraspecific trait variation on trait-based niches and performance of legumes in plant mixtures. Plos One 10, e0119786.

Roscher, C., Thein, S., Schmid, B., Scherer-Lorenzen, M., 2008. Complementary nitrogen use among potentially dominant species in a biodiversity experiment varies between two years. J. Ecol. 96, 477–488.

Rottstock, T., Kummer, V., Fischer, M., Joshi, J., 2017. Rapid transgenerational effects in *Knautia arvensis* in response to plant community diversity. J. Ecol. 105, 714–725.

Roughgarden, J., 1974. Niche width: biogeographic patterns among Anolis lizard populations. Am. Nat. 108, 429–442.

Schmid, B., 1985. Clonal Growth in Grassland Perennials: III. Genetic Variation and Plasticity Between and Within Populations of Bellis Perennis and Prunella Vulgaris. J. Ecol. 73, 819–830.

Schnitzer, S.A., Klironomos, J.N., HilleRisLambers, J., Kinkel, L.L., Reich, P.B., Xiao, K., Rillig, M.C., Sikes, B.A., Callaway, R.M., Mangan, S.A., others, 2011. Soil microbes drive the classic plant diversity-productivity pattern. Ecology 92, 296–303.

Schoener, T.W., Gorman, G.C., 1968. Some niche differences in three lesser antillean lizards of the genos Anolis. Ecology 49, 819–830.

Siefert, A., et al., 2015. A global meta-analysis of the relative extent of intraspecific trait variation in plant communities. Ecol. Lett. 18, 1406–1419.

Soliveres, S. et al., 2016. Biodiversity at multiple trophic levels is needed for ecosystem multifunctionality. Nature 536, 456–459.

Spehn, E.M., Joshi, J., Schmid, B., Diemer, M., Korner, C., 2000. Above-Ground Resource Use Increases with Plant Species Richness in Experimental Grassland Ecosystems. Funct. Ecol. 14, 326–337.

Sterck, F., Markesteijn, L., Schieving, F., Poorter, L., 2011. Functional traits determine trade-offs and niches in a tropical forest community. Proc. Natl. Acad. Sci. 108, 20627–20632.

Thorpe, A.S., Aschehoug, E.T., Atwater, D.Z., Callaway, R.M., 2011. Plant interactions and evolution. J. Ecol. 99, 729–740.

Tilman, D., Reich, P.B., Knops, J., Wedin, D., Mielke, T., Lehman, C., 2001. Diversity and productivity in a long-term grassland experiment. Science 294, 843–845.

Turcotte, M.M., Levine, J.M., 2016. Phenotypic Plasticity and Species Coexistence. Trends Ecol. Evol. 31, 808–813.

van Ruijven, J., Berendse, F., 2005. Diversity-Productivity relationships: initial effects, long-term patterns, and underlying mechanisms. Proc. Natl. Acad. Sci. U. S. A. 102, 695–700.

van Valen, L., 1965. Morphological variation and width of ecological niche. Am. Nat. 99, 377–390.

van Moorsel, S. J., Hahl, T., Wagg, C., De Deyn, G. B., Flynn, D.F.B., Zuppinger-Dingley, D., Schmid, B., 2017. Community evolution increases plant productivity at low diversity. bioRxiv 111617.

Vermeulen, P.J., Anten, N.P.R., Schieving, F., Werger, M.J.A., During, H.J., 2008. Height convergence in response to neighbour growth: genotypic differences in the stoloniferous plant *Potentilla reptans*. New Phytol. 177, 688–697.

Violle, C., Enquist, B.J., McGill, B.J., Jiang, L., Albert, C.H., Hulshof, C., Jung, V., Messier, J., 2012. The return of the variance: intraspecific variability in community ecology. Trends Ecol. Evol. 27, 244–252.

Violle, C., Navas, M.-L., Vile, D., Kazakou, E., Fortunel, C., Hummel, I., Garnier, E., 2007. Let the concept of trait be functional! Oikos 116, 882–892.

von Felten, S., Hector, A., Buchmann, N., Niklaus, P.A., Schmid, B., Scherer-Lorenzen, M., 2009. Belowground nitrogen partitioning in experimental grassland plant communities of varying species richness. Ecology 90, 1389–1399.

Wacker, L., Baudois, O., Eichenberger-Glinz, S., Schmid, B., 2009. Effects of plant species richness on stand structure and productivity. J. Plant Ecol. 2, 95–106.

White, J.W., Montes-R, C., 2005. Variation in parameters related to leaf thickness in common bean (Phaseolus vulgaris L.). Field Crops Res. 91, 7–21.

Williams, L.J., Paquette, A., Cavender-Bares, J., Messier, C., Reich, P.B., 2017. Spatial complementarity in tree crowns explains overyielding in species mixtures. Nat. Ecol. Evol. 1, 63.

Zuppinger-Dingley, D., Schmid, B., Petermann, J.S., Yadav, V., De Deyn, G.B., Flynn, D.F.B., 2014. Selection for niche differentiation in plant communities increases biodiversity effects. Nature 515, 108–111.

